# MinION Nanopore Sequencing of Multiple Displacement Amplified Mycobacteria DNA Direct from Sputum

**DOI:** 10.1101/490417

**Authors:** Sophie George, Yifei Xu, Nicholas Sanderson, Alasdair TM Hubbard, David T. Griffiths, Marcus Morgan, Louise Pankhurst, Sarah J. Hoosdally, Dona Foster, Samantha Thulborn, Esther Robinson, E. Grace Smith, Priti Rathod, A. Sarah Walker, Timothy E. A. Peto, Derrick W. Crook, Kate E. Dingle

**Affiliations:** Nuffield Department of Clinical Medicine, John Radcliffe Hospital, Oxford University, UK; National Institute for Health Research (NIHR) Oxford Biomedical Research Centre, John Radcliffe Hospital, Oxford, UK; NIHR Oxford Health Protection Research Unit in Healthcare Associated Infection and Antimicrobial Resistance at Oxford University in partnership with Public Health England, Oxford, UK; Microbiology Department, Oxford University Hospitals NHS Trust, Oxford, UK; Respiratory Medicine Unit, Nuffield Department of Medicine, John Radcliffe Hospital, University of Oxford, UK; PHE National Mycobacteria Reference Service - North and Central, Birmingham Public Health Laboratory, UK

## Abstract

Sequencing of pathogen DNA directly from clinical samples offers the possibilities of rapid diagnosis, faster antimicrobial resistance prediction and enhanced outbreak investigation. The approach is especially advantageous for infections caused by species which grow very slowly in culture, such as *Mycobacteria tuberculosis*. Since the pathogen of interest may represent as little as 0.01% of the total DNA, enrichment of the input material for target sequences by specific amplification and, or depletion of non-target DNA (human, other bacteria) is essential for success. Here, we investigated the potential of isothermal multiple displacement amplification by Phi29 polymerase. We directed the amplification reaction towards Mycobacteria DNA in sputum samples by exploiting in our oligonucleotide primer design, their high GC content (approximately 65%) relative to human DNA. Amplified DNA was then sequenced using the Oxford Nanopore Technology MinION. In addition, a model system comprising standardised ‘mock clinical samples’ was designed. Pooled infection negative human sputum samples were spiked with enumerated *Mycobacterium bovis* (BCG) Pasteur strain at concentrations spanning the typical range at which *Mycobacterium tuberculosis* is found in human sputum samples (10^6^ - 10^1^ BCG cells/ml). To assess the amount of BCG sequence enrichment achieved, sample DNA was sequenced both before, and after amplification. Reads from amplified samples, which mapped to a BCG reference genome, comprised short repeated sequences - apparently transcribed multiple times from the same fragment of BCG DNA. Therefore post-amplification, the samples were enriched for BCG sequences relative to unamplified sequences (8,101 BCG reference mapped reads, increasing to 28,617 at 10^6^ BCG cells/ml sample), but BCG genome coverage declined markedly (for example 89.4% to 4.1%). In summary, the use of standardised mock clinical samples allowed direct comparison of data from different Mycobacteria enrichment experiments and sequencing runs. However, optimal conditions for multiple displacement amplification of minority Mycobacteria DNAs, remain to be identified.

## INTRODUCTION

The World Health Organization (WHO) estimates that in 2016, *Mycobacterium tuberculosis* complex caused 6.3 million new TB cases and 1.67 million deaths worldwide (including 374,000 among HIV-positive people) [1]. In addition, *Mycobacterium abscessus, M. avium* complex, *M. kansasii, M. malmoense*, and *M. xenopi* are currently the most clinically important of the >160 known non-tuberculous mycobacteria (NTM) species [2-4]. Correct diagnosis and antimicrobial resistance determination are essential to ensure appropriate treatment of Mycobacteria infections. However, when based on growth in culture, this may take up to 80 days from initial presentation, increasing the risk of poor clinical outcomes and failure to identify and control transmission.

Routine whole genome sequencing of Mycobacteria using Illumina MiSeq has accelerated laboratory diagnosis of Mycobacteria by Public Health England (PHE) [5, 6]. Samples are cultured until positive, which usually occurs within 1-2 weeks if the sample is smear positive (but up to 5-6 weeks if bacterial load is low), then total DNA is extracted and sequenced using the Illumina platform [5, 7]. WGS diagnostics can be completed in a median of 9 days (IQR 6-10) [5]. Antimicrobial resistance predictions are based on nucleotide sequence data [8], and phylogenetic analyses identify transmission events and outbreaks [9, 10]. The information gained from WGS methods is typically available to the clinician within three to four weeks of the sample being taken. Further savings in cost and time could potentially be achieved by determining Mycobacteria genome sequences from DNA extracted directly from clinical samples, thus eliminating the need for culture altogether.

Whole genome sequencing of pathogens direct from clinical samples is technically challenging. Samples vary in terms of volume, numbers of human and bacterial cells and the concentration of target organisms. Mycobacteria DNA, for example, can represent as little as 0.01% of the total DNA extracted from sputum [11]. Small scale studies employing direct from sample sequencing have reported 0.002 - 0.7% sequence coverage of the *M. tuberculosis* genome (using differential lysis and a DNA extraction kit) [12], and up to 90% genome coverage with 20x depth (20/24 samples), (using the SureSelect target enrichment method, Agilent, USA) [13, 14]. The study by Brown et al. [13] predicted both Mycobacteria species and antibiotic susceptibility, but the cost ($350 per sample) and duration of the protocol (2 to 3 days) could prevent its use. The ideal ‘direct from sample’ methodology would be simple, low cost and portable, to facilitate use in remote, low-income settings where the burden of infection is greatest, and provision of clinical diagnostic services and treatment is severely limited.

Potential advantages of adopting the Nanopore sequencing platform (Oxford Nanopore Technology, ONT, Oxford, UK) include the possibility of increased read lengths [15, 16] and consequent improved *de novo* assemblies, avoiding the need for a reference genome [16]. The accuracy of DNA sequences obtained using the Nanopore platform is constantly improving; 99.9% can be achieved when Nanopolish is used to improve consensus accuracy [17].

Enrichment of target pathogen sequences within total extracted DNA is a prerequisite for direct-from-sample sequencing. The technique of isothermal multiple displacement amplification (MDA) using Phi29 DNA polymerase [18] shows promise, since μg quantities of DNA can be generated from minimal template (1-10ng) [19-21]. In the present study, we investigated the possibility of biasing MDA towards Mycobacteria DNA in sputum samples, prior to sequencing the DNA directly using the Oxford Nanopore Technology MinION.

## MATERIALS AND METHODS

### Standardised Samples for Method Development

A model system comprising standardised ‘mock clinical samples’ was designed. Pooled infection negative human sputum samples were spiked with enumerated “Bacille de Calmette et Guérin” *Mycobacterium bovis* (BCG) Pasteur strain (attenuated derivative of *Mycobacterium bovis* [22]) at known concentrations. This allowed the results of different experiments to be compared.

### BCG Culture and Enumeration

BCG Mycobacteria Growth Incubator Tube (MGIT) culture conditions were optimised with a two-step process facilitating the growth of single, rather than clusters or ‘flakes’ of BCG cells. Firstly, freshly prepared MGIT culture tubes (Becton Dickinson, Wokingham, UK) were inoculated sparsely with 10 μl BCG frozen stock. After 30 days incubation at 37°C, the cultures were vortexed vigorously. ‘Flakes’ comprising large numbers of BCG cells were allowed to settle for 10 minutes. Fresh MGIT tubes were prepared, with the addition of Tween 80 (Acros Organics, Geel, Belgium) (0.5% final concentration) to encourage BCG growth as single cells [23]. These fresh tubes were inoculated using 200 μl ‘settled’ BCG culture. After 18 days incubation at 37°C, BCG cells were harvested and counted as follows.

The BCG culture was vortexed vigorously and 1 ml was removed. ‘De-clumped’ BCG cells were pelleted by centrifugation for 10 minutes (13,000 rpm), then the pellet was resuspended in 100 μl crystal violet stain (Pro Lab Diagnostics, Birkenhead, UK). Cells were counted using a Petroff Hausser counting chamber (Hausser Scientific, Horsham, PA, USA) for bacteria enumeration. After counting, the enumerated BCG stock (in MGIT culture fluid) was stored at −20 °C in 1 ml aliquots until required. At this point, a 10 fold dilution series of BCG cells was made in phosphate buffered saline.

Mock clinical samples were prepared by pooling ten anonymised infection negative sputum samples (from asthmatic patients) ((see research ethics statement below). Pooled sputum was liquefied by treatment with an equal volume of freshly prepared working strength Sputasol (Oxoid, Thermo Scientific, Paisley, UK). The sputum was incubated at 37°C with occasional vortexing, until liquefaction was complete. 1ml aliquots of the negative sputum samples were spiked with cells from the BCG titration.

### Research Ethics Statement

The protocol for this study was approved by London – Queen Square Research Ethics Committee (17/LO/1420). Human samples were collected under approval of East Midlands Research Ethics Committee (08/H0406/189) and all subjects gave written informed consent in accordance with the Declaration of Helsinki.

### DNA Extraction directly from Mock Clinical Samples

Each sample underwent a saline wash to remove extraneous human DNA. After centrifugation at 13,200 rpm for 15 minutes the supernatant was discarded, then the pellet was resuspended in 1 ml sterile phosphate buffered saline and centrifugation repeated. The pellet was transferred in 100 μl molecular grade H_2_O to a 0.5 ml plain skirted tube (STARLAB, Hamburg, Germany) containing 0.8 g of aliquoted 0.1 mm silica beads (lysing matrix B, MP Biomedicals, Santa Ana, USA). The mixture underwent bead beating (3×40s, 3 minute interval, 6.0 m/s) on a Fast Prep-24 machine (MP Biomedicals) followed by centrifugation at 13,200 rpm for 10 minutes. DNA was recovered from 50 μl supernatant using 1.8x volume magnetic AMPure XP beads (Beckman Coulter, High Wycombe, UK). After vortexing for 20 seconds, and magnetic separation for 10 minutes, the supernatant above the beads was replaced with 200μl of 80% EtOH. This was removed after 1 minute and the wash step repeated, after which the beads were air dried for 10 minutes. DNA was eluted in 26μl of 1 x TE buffer (pH8, Sigma Aldrich, Dorset, UK). DNA concentration, integrity and fragment size were measured by Qubit Fluorometer (Rugby, UK) and TapeStation (Stockport, UK) respectively.

### Multiple Displacement Amplification by Phi29 DNA polymerase

DNA (1 ng/10 μl) extracted from mock clinical samples was denatured by incubation for 3 minutes at 96 °C, then transferred to ice, where the rest of the MDA reaction was assembled. The final 20 μl reaction comprised 1x phi29 DNA polymerase reaction buffer (New England BioLabs, Hitchin, UK), 0.1 mg/ml BSA, 5 mM dNTPs, 10 μM oligonucleotide primers (see below) with a modified 3’-terminal endonuclease resistant phosphorothioate bond, 5 mM MgCl_2_, and 2 μl Phi29 DNA polymerase (20 units) (New England Biolabs). Two alternative primers were tested; ‘random’ hexamers containing 65% GC, or ‘most frequent 10mers’ based on the most frequent 10 bp sequence repeats identified in Mycobacteria genome (S1 Table). Incubation was at 30 °C for 16 hours. Amplified DNA was purified using AMPure XP beads, quantitated, and 1 μg was digested to remove branched structures using 1 μl T7 Endonuclease I (New England BioLabs, Hitchin, UK) in 20 μl reaction volume at 37°C for 1 hour, followed by a second AMPure XP bead purification step.

### Oxford Nanopore Technology (ONT) Sequencing Library Preparation and Sequencing

Digested DNA (700-900 ng/μl) was prepared for ONT sequencing according to the manufacturer’s 1D Native barcoding genomic DNA protocol using SQK-LSK108 and EXP-NBD103 kits (Oxford Nanopore Technology, Oxford, UK). Each sequencing library comprised seven barcoded DNA samples and was sequenced using MinION R9.4 SpotON flow cells for 48 hours.

### Bioinformatics

MinION reads were basecalled using Albacore v2.0.2 (Oxford Nanopore Technology, Oxford, UK). We used Porechop (v0.2.2, https://github.com/rrwick/Porechop) to perform stringent barcode demultiplexing of the sequencing data. Porechop confirms the presence of the barcode sequence at both the start and end of each read; reads were acceptable only if the same barcode was found at both ends, otherwise the read was discarded. This level of stringency was achieved by setting the “require_two_barcodes” option in Porechop and setting the threshold for the barcode score at 60. The basic statistics of the sequencing data were reported using NanoPack [24]. Then, reads from each sample were mapped to the BCG reference sequence (GenBank AM408590; the 16S rRNA region {1498360, 1499896} was masked) using Minimap2 [25]. Integrative Genomics Viewer [26] was used to view the resulting alignment profiles. The number of reads (i) mapped to the BCG reference, (ii) fitting the definition of supplementary alignments (as below), and (iii) alignment length were analysed using Pysam (https://github.com/pysam-developers/pysam). Repeated BCG-derived sequences were found within the contiguous sequence of certain individual reads. These reads therefore could not be aligned linearly to the BCG reference. One of the linear BCG repeats within such reads was referred to as the “representative alignment” and the additional repeats were referred to as “supplementary alignment(s)”. ‘Supplementary alignments’ were considered to be present if the start and end of their BCG alignment positions occurred within 10 bp of the representative alignment, ie up to 10 bp could occur between the repeated sequences. The histogram for the alignment length against the read length and the number of repeats in each read was plotted by using ggplot2 implemented in R (https://www.r-project.org/).

## RESULTS

### Experimental Design

The range of Mycobacteria cell concentrations typically found in *Mycobacterium tuberculosis*-positive sputum [27] were represented in standardised mock clinical samples, the BCG dilution series ranging from 10^6^ to 10^1^ BCG cells per ml liquefied sputum. Total DNA was extracted from these sputum samples and a negative sputum control. The DNA was sequenced on the R9.4 flow cell, to establish the proportions of BCG and other DNAs present (Table 1: no amplification). This extracted DNA then formed the template for testing molecular methods based on differently primed phi29 amplification reactions, aiming to selectively enrich for BCG DNA within the total (Table 1: amplification).

**Table 1:** Comparison of sequence data obtained for BCG-spiked sputum samples; with and without prior amplification. Number of reads, mapped reads, genome coverage, supplementary alignment, and mean alignment length for un-amplified samples and phi29 amplified samples (different primers) at the concentrations of BCG cells shown. Supplementary alignments occurred when the contiguous sequence of an individual read could not be aligned linearly to the reference sequence. Thus, one of the linear alignments in a repeat-containing read is referred as the “representative alignment” and the others (repeats of this sequence) are referred as “supplementary alignment(s)”. * ratio not given when the number of mapped read is less than 10.

| Amplification | BCG concentration (cells/ml) | Number of reads | Mapped reads | Genome coverage (%) | Supplementary alignment (ratio to mapped reads*) | Mean Alignment length |
| --- | --- | --- | --- | --- | --- | --- |
| None | 10<sup>6</sup> | 71,029 | 8,068 | 89.4 | 125 (0.0) | 1305 |
|  | 10<sup>5</sup> | 38,752 | 491 | 13.3 | 2 (0.0) | 1276 |
|  | 10<sup>4</sup> | 9,543 | 11 | 0.3 | 0 (0.0) | 1106 |
|  | 10<sup>3</sup> | 98,845 | 20 | 0.5 | 0 (0.0) | 1047 |
|  | 10<sup>2</sup> | 96,293 | 2 | 0.2 | 1 | 2402 |
|  | 10<sup>1</sup> | 32,743 | 0 | 0.1 | 0 |  |
|  | 0 | 70,929 | 4 | 0.2 | 0 | 1168 |
| Phi29 65% GC primers | 10<sup>6</sup> | 144,331 | 28,617 | 4.1 | 12,0415 (4.2) | 205 |
|  | 10<sup>5</sup> | 126,490 | 7,359 | 1.2 | 27602 (3.8) | 214 |
|  | 10<sup>4</sup> | 58,851 | 84 | 0.1 | 495 (5.9) | 171 |
|  | 10<sup>3</sup> | 76,870 | 10 | 0.1 | 28 (2.8) | 210 |
|  | 10<sup>2</sup> | 105,612 | 0 | 0.0 | 0 |  |
|  | 10<sup>1</sup> | 92,841 | 1 | 0.0 | 1 | 162 |
|  | 0 | 139,173 | 1 | 0.0 | 2 | 309 |
| Phi29 MF 10mer primers | 10<sup>6</sup> | 201,038 | 31,893 | 4.6 | 54,651 (1.7) | 181 |
|  | 10<sup>5</sup> | 112,595 | 7,405 | 1.6 | 14,759 (2.0) | 322 |
|  | 10<sup>4</sup> | 148,550 | 2 | 0.0 | 0 | 342 |
|  | 10<sup>3</sup> | 148,612 | 4 | 0.0 | 1 | 116 |
|  | 10<sup>2</sup> | 186,505 | 1 | 0.0 | 1 | 145 |
|  | 10<sup>1</sup> | 91,657 | 0 | 0.0 | 0 |  |
|  | 0 | 275,190 | 2 | 0.0 | 1 | 102 |

### Selective Enrichment of Mycobacteria sequences using Phi29 DNA polymerase

The high GC content of the Mycobacteria genome (for *M. tuberculosis* 65.6% GC) [28] relative to most of the human genome (<50% GC for ∼92% of the genome and 50-60% GC in ∼7% genome [29] was exploited in our experimental design. MDA was primed using 65% GC biased ‘random’ hexamers, or MF (most frequent) 10mer primers (S1 Table). An unamplified DNA control was included at each sample concentration. Amplification products (or control DNA) (200ng) were sequenced using the Oxford Nanopore Technologies (ONT) MinION R9.4 platform (see S2 Table for summary of sample DNA concentration, amplification, sequencing library preparation, and statistics of sequencing data).

For the unamplified BCG-spiked samples, the percentage of total DNA reads which mapped to the BCG reference increased with increasing BCG cell concentration (Table 1). For example, at 10^6^ cells/ml, 11.4% of total reads mapped to the BCG reference and 89.4% of the BCG genome was covered. The comparable results for Phi29 amplified (65% GC hexamer primed) extracted DNA showed an increased proportion of BCG reads within the total - 19.8% of the total reads. However, BCG genome coverage was much lower, at 4.1% (Table 1 upper panel).

The BCG reference-mapped alignment profiles of samples which had undergone Phi29 amplification (65% GC hexamer primed) contained a large number of ‘supplementary alignments’ (as defined in Materials and Methods) (Table 1). This repeat feature was virtually absent in the sequence data from non-Phi29 amplified samples. The ratios between Phi29 amplified sequences forming representative and supplementary alignments were 4.2 (at 10^6^ BCG cells/ml), 3.8 (10^5^), 5.9 (10^4^), and 2.8 (10^3^), respectively (Table 1). The mean BCG alignment length (about 200 nucleotides) in the 65% GC hexamer-primed Phi29 amplified sequences was considerably shorter than that of the standard, unamplified sequence data.

### Explanation for biased Phi29 amplification

To explain these findings, we hypothesised that Phi29 DNA Polymerase enriched the samples in terms of their total BCG derived DNA content, but amplified short sequences to high depths, ie the reads that mapped to the BCG reference in the Phi29 amplified data comprised short repeated sequences potentially transcribed from the same fragment of DNA.

Visualization of the mapping profile revealed that BCG-like reads were split into multiple small fragments which each mapped to same region of the reference genome (Fig 1). More than 67% of the reads (at 10^6^ BCG cells/ml) which mapped to the BCG reference comprised repeats. Two, three, four, and five repeats were observed in 15.7%, 11.3%, 8.4%, and 6.3% of the reads, respectively.

**Fig 1.**
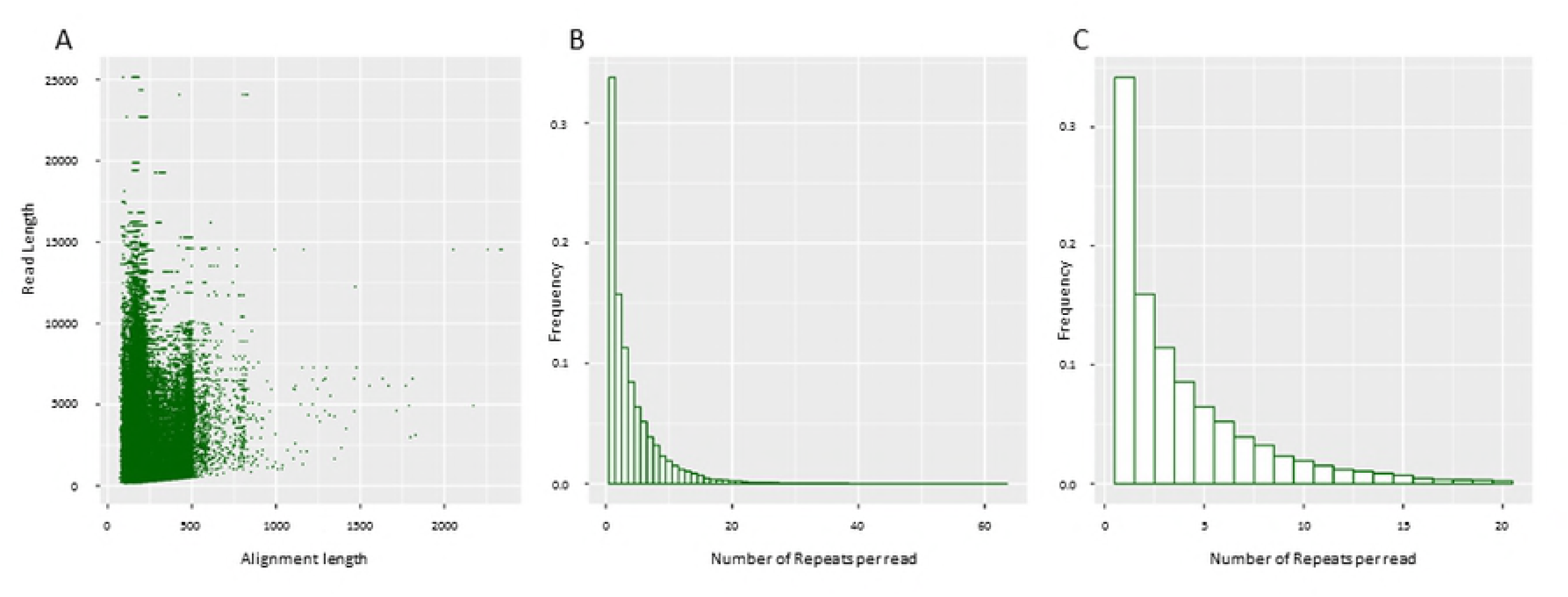
Analysis of reference BCG mapping profile for Phi29 amplified sequence data at 106 cells/ml BCG. (A) Plot of alignment length against read length. (B) Histogram showing number of repeats in each read that mapped to BCG reference. (C) The same histogram as (B) but focusing only on the number of repeats within the range from 1 to 20 per read.

The overall GC content of the Phi29 ‘65% GC hexamer-primed’ amplified sequences was very close to the BCG average of 65.6%. The BCG sequences amplified from the 10^5^ and 10^6^spiked samples contained GC 65.5% (within 1.2% genome coverage) and 65.7% (in 4.1% genome coverage) respectively, compared to 65.6% across the whole of the genome. This suggests there was no amplification bias towards BCG sequences which have GC content away from the mean, and that the repeatedly amplified BCG sequences were not unusual in terms of their overall GC content. The distribution of the most abundant Phi29 amplification products across the sequence of the BCG reference genome also indicated that obvious amplification hot spots were absent (Fig 2).

**Fig 2.**
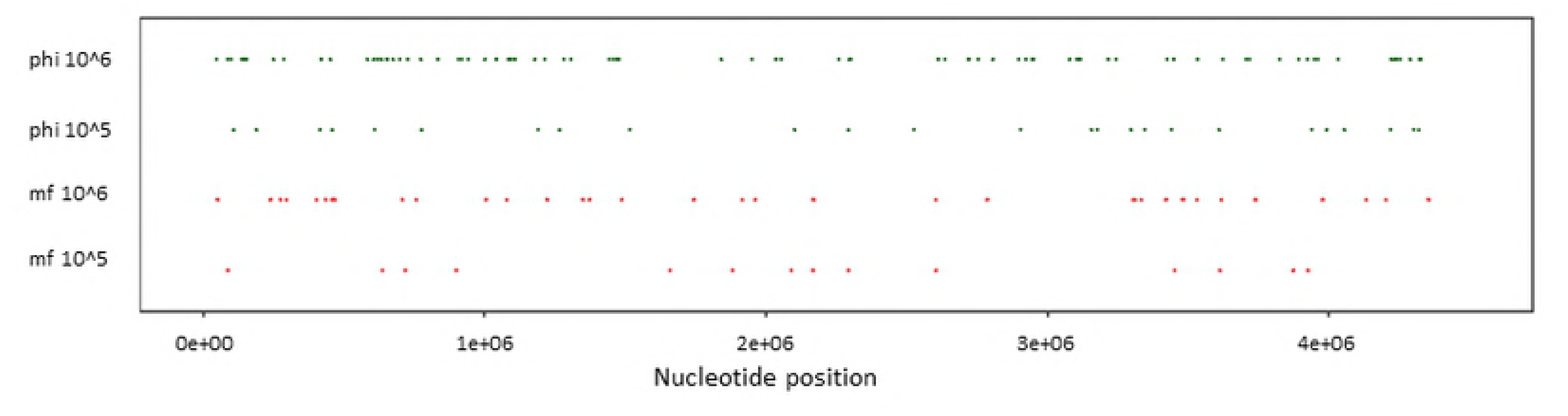
Distribution of most abundant Phi29 amplification products relative to the BCG reference genome indicates the absence of obvious amplification hot spots. Comparison of the regions of the BCG genome amplified by Phi29 primed either using 65% GC hexamers (phi) or most frequent 10mers (mf) at 10^6 and 10^5 BCG cells/ml. The top 10% nucleotide positions with highest depth of mapping coverage are shown.

### BCG Reads detected in the Negative Control/Low Concentration Spikes

The negative control sample (negative sputum with zero BCG cells added) contained a small number of reads (less than five) which mapped to the BCG reference (Table 1). This also occurred in sputum samples spiked at low BCG concentrations (10^2^ and 10^1^ cells/ml) in both the phi29 amplified and unamplified control sequence data. All samples had been sequenced while ‘multiplexed’ – the addition of a barcode sequence to each sample during library preparation allowed ‘de-multiplexing’ to be performed bioinformatically after sequencing. Despite the fact that we implemented stringent bioinformatic barcode removal for de-multiplexing, which successfully removed most of this cross-contamination, a low level remained. This issue was confirmed to be bioinformatics-based, when samples were run of single flow cells (not multiplexed).

## DISCUSSION

Multiple Displacement Amplification of DNA by Phi29 polymerase is an attractive choice for experiments aiming to generating large quantities of DNA (≥μg) from very small (≤ng) amounts under isothermal conditions [18]. Advantages include a low error rate due to 3’,5’-exonuclease ‘proofreading’ activity (error rate ∼9.5 x 10^-6^), the capacity to synthesise DNA molecules >70kb long and the possibility of virtually whole genome amplification [19, 30-32]. Relative to PCR-based methods, more DNA is amplified by at least an order of magnitude, and good genome coverage and reduced amplification bias of genomic DNA from human cells has been reported [33]. Long DNA fragments provide ideal input for the Nanopore MinION sequencing platform, which in turn generates long reads offering the possibility of *de novo*, rather than reference based genome assemblies.

MDA has also shown promise for the accurate and unbiased amplification of whole bacterial genomes from uncultivable, or slow growing species, and even ‘single cell genomics’ [34, 35]. MDA with random hexamers has been used to amplify *Xylella fastidiosa* (Gram negative plant pathogen, 52% CG content) DNA directly from approximately 1000 target cells, yielding over 4 μg of high molecular weight DNA and achieving uniform genome coverage relative to unamplified DNA [34]. Coverage of *Coxiella burnetii* (fastidious obligate intracellular pathogen, 42.5% GC) was similarly representative, as assessed by PCR [36]. Work in our laboratory aiming to sequence Mycobacteria directly from sputum samples has previously used 3% NaOH (Nac-Pac Red; Alpha-Tec Systems, Vancouver, WA, USA) to deplete sputum of non-Mycobacteria, together with a ‘Molysis kit’ (Molzym Life Science, Bremen, Germany) to reduce human DNA contamination [11]. An important issue arising from these pre-treatments is that for most samples, insufficient DNA remains for direct sequencing using the Nanopore MinION. Here, we aimed to investigate the possibility of eliminating the need for such pre-treatment, while amplifying microgram quantities of DNA enriched for Mycobacteria sequences.

Our experiments employed 65% GC biased hexamers to favour amplification of the BCG genome (65% GC content) relative to the human genome (CG content <50% for ∼92% of the genome and 50-60% CG for ∼7% genome [29]). This achieved two to five fold enrichment for BCG sequences (Table 1) but at the expense of genome coverage (for example 89.4% genome coverage decreased to 4.1% at the 10^6^ BCG spike concentration, Table 1). Post amplification, certain regions of the genome were covered at extremely high depth. The reason for the high coverage in certain regions, but not others is unknown. It may be unrelated to the mean GC content of these sequences, because this was the same within amplified sequences as the mean for the whole genome. The absence of obvious amplification hotspots conserved between experiments (Fig 2) suggests regions of high coverage may occur stochastically.

The difficulty of amplifying GC rich sequences from a complex mixture by MDA has been reported previously [37]; species with the highest GC content underwent significantly less amplification from an environmental (soil) sample compared to low GC bacteria. Yilmaz et al. [38] evaluated three different commercially available kits, including NEB Phi29 used in our study. They also observed amplification bias against high (G+C)-content templates in bacteria amplified from sludge and compost communities. Our use of 65% GC biased hexamers (also MF10mers Table 1) with the NEB Phi29 polymerase was insufficient to achieve unbiased amplification of the GC rich BCG genome. Similar bias against GC rich sequences has been observed previously [39]; MDA of DNA extracted from tumour samples reproducibly distorted gene dosage representation in the amplified DNA, reflecting the GC content of different regions of the template. Also, a study of copy number variants within the human genome created hundreds of potentially confounding MDA artefacts that could obscure authentic copy number variants, which were reproducible and influenced by GC content [40]. There is also evidence of stochastic effects originating from the amplification of very low amounts of genomic template from a single bacterium [41] - locus representation values ranged from 0.1% to 1,211%.

The reason MDA is biased against GC rich templates is unclear, but it could reflect the higher melting temperature of GC rich DNA relative to AT rich sequences. In addition to the conditions described above, we tested reaction conditions which are known to alleviate GC-melting related issues in PCR, by effectively reducing the melting temperature of the DNA (PCR additives Q-solution and DMSO), as well as increasing the incubation temperature to 35 °C and 40 °C. A novel thermostable mutant of Phi29 polymerase, designated WGA-X (Thermofisher) has been described which amplifies DNA at 45°C, [42] and offers improved amplification of high GC content templates, but it was not commercially available. We also tested the Phi29-polymerase based Qiagen REPLI-g kit (data not shown) because it uses alkaline DNA denaturation to improve the uniformity of DNA denaturation while minimising DNA fragmentation or generation of abasic sites (relative to heat denaturation), and because it’s been reported to work at 40°C [43]. This kit was also tested by Yilmaz et al. [38] and it performed best with respect to GC bias. Unfortunately, none of these modifications improved the genome coverage achieved in our study. Further experiments with shorter amplification incubation times were also performed, with the aim of potentially reducing the amplification bias, but these were unsuccessful.

The challenges of amplifying a minority, GC rich target DNA from within a complex mixture remain. Here, establishing mock clinical samples containing defined numbers of BCG cells represented a key step forward, because the data from different method development experiments could be compared. This material is proving invaluable in further work aiming to optimise ‘direct from sample’ Mycobacteria genome sequencing. In conclusion, optimal conditions under which Phi29 polymerase might be directly amplify minority GC rich templates without bias, remain to be identified.

## ACKNOWLEDGEMENTS

The views expressed in this publication are those of the authors and not necessarily those of the NHS, NIHR, the Department of Health or Public Health England.

DWC and TEAP are NIHR senior investigators.

## AUTHOR CONTRIBUTIONS

**Conceptualization and Methodology:** Sophie George, Yifei Xu, Alasdair TM Hubbard, Louise Pankhurst, Sarah J Hoosdally, Dona Foster, A. Sarah Walker, Timothy EA Peto, Derrick W Crook, Kate. E Dingle.

**Resources:** Samantha Thulborn, Esther Robinson, E Grace Smith, Priti Rahood, Marcus Morgan.

**Investigation:** Sophie George, Yifei Xu, Alasdair TM Hubbard, David T Griffiths, Marcus Morgan, Kate E Dingle.

**Formal Analysis**: Yifei Xu, Nicholas Sanderson, Timothy EA Peto.

**Software**: Yifei Xu, Nicholas Sanderson.

**Visualisation:** Sophie George, Yifei Xu, Timothy EA Peto, Kate E Dingle.

**Writing – Original Draft Preparation**: Kate E Dingle.

**Writing – Review & Editing**: Sophie George, Yifei Xu, Alasdair TM Hubbard, David T Griffiths, Marcus Morgan, Louise Pankhurst, Sarah J Hoosdally, Dona Foster, Samantha Thulborn, Esther Robinson, E Grace Smith, Priti Rahood, A. Sarah Walker, Timothy EA Peto, Derrick W Crook, Kate. E Dingle.

## CONFLICT OF INTERESTS

The authors declare no conflicts of interest.

